# Enriched G-quadruplexes on the *Drosophila* Male X Chromosome Function as Insulators of Dosage Compensation Complex

**DOI:** 10.1101/656538

**Authors:** Sheng-Hu Qian, Lu Chen, Zhen-Xia Chen

**Affiliations:** Hubei Key Laboratory of Agricultural Bioinformatics, College of Life Science and Technology, Huazhong Agricultural University, Wuhan, Hubei 430070, PR China

## Abstract

The evolution of sex chromosomes has resulted in half X chromosome dosage in males as females. Dosage compensation, or the two-fold upregulation in males, was thus evolved to balance the gene expression between sexes. However, the step-wise evolutionary trajectory of dosage compensation during Y chromosome degeneration is still unclear. Here, we show that the specific structured elements G-quadruplexes (G4s) are enriched on the X chromosome in *Drosophila melanogaster*. Meanwhile, on the X chromosome, the G4s are underrepresented on the H4K16 acetylated regions and the binding sites of dosage compensation complex male-specific lethal (MSL) complex. Peaks of G4 density and potential are observed at the flanking regions of MSL binding sites, suggesting G4s act as insulators to precisely up-regulate certain regions in males. Thus, G4s may be involved in the evolution of dosage compensation process through fine-tuning one-dose proto-X chromosome regions around MSL binding sites during the gradual Y chromosome degeneration.

**One Sentence Summary:** G-quadruplexes act as insulators to precisely up-regulate X chromosome in males.

## Main Text

Dosage compensation in flies represents a prime example of fine-tuning gene expression of a whole chromosome. It balanced the gene expression between females (XX) and males (XY) through the twofold upregulation of X chromosome in males mediated by male-specific lethal (MSL) complex (*1, 2*). MSL complex lands on genomic regions called High-Affinity Sites (HAS), and then spreads to acetylate histone H4 lysine 16 (H4K16ac) across the entire X chromosome (*3*). Recent advances reveal that the complex interplay among the epigenetic landscape, transcription, and chromosome conformation achieves male-specific X chromosome upregulation (*4, 5*). However, the detailed mechanisms underlying dosage compensation are still obscure.

G-quadruplexes (G4s), the four-stranded helical structures formed by guanine-rich DNA or RNA (Fig. 1A), impose a substantial mutational burden on the genome (*6–8*). However, a growing body of evidence implicating G4s as regulators of gene expression (*9, 10*). To explore the functionality of G4s, we systematically searched for potential G4s (pG4s) in 20 *Drosophila* species, and identified 43,244-120,764 pG4s across their genomes (Fig. S1). Specifically, 46,435 pG4s were found in *D. melanogaster* (1.12% of the genome), including 15,663 pG4s in intergenic regions, and even 8.5 times as high as in randomly shuffled genome (Fig. S2). The density of pG4s in intergenic regions was higher than that in genic regions (Fig. 1B). Besides, although GC contents of sliding windows across the genome were positively correlated with the pG4 number (Fig. S3; r = 0.359, P < 2.2 × 10^-16^), the GC contents of intergenic regions were lower than genic regions (Fig. S4), suggesting a maintenance of intergenic pG4s as functional elements under selection. Alternatively, pG4s may be neutral or nearly neutral, and more pG4 were maintained in intergenic regions merely due to less selective constraints.

**Fig. 1.**
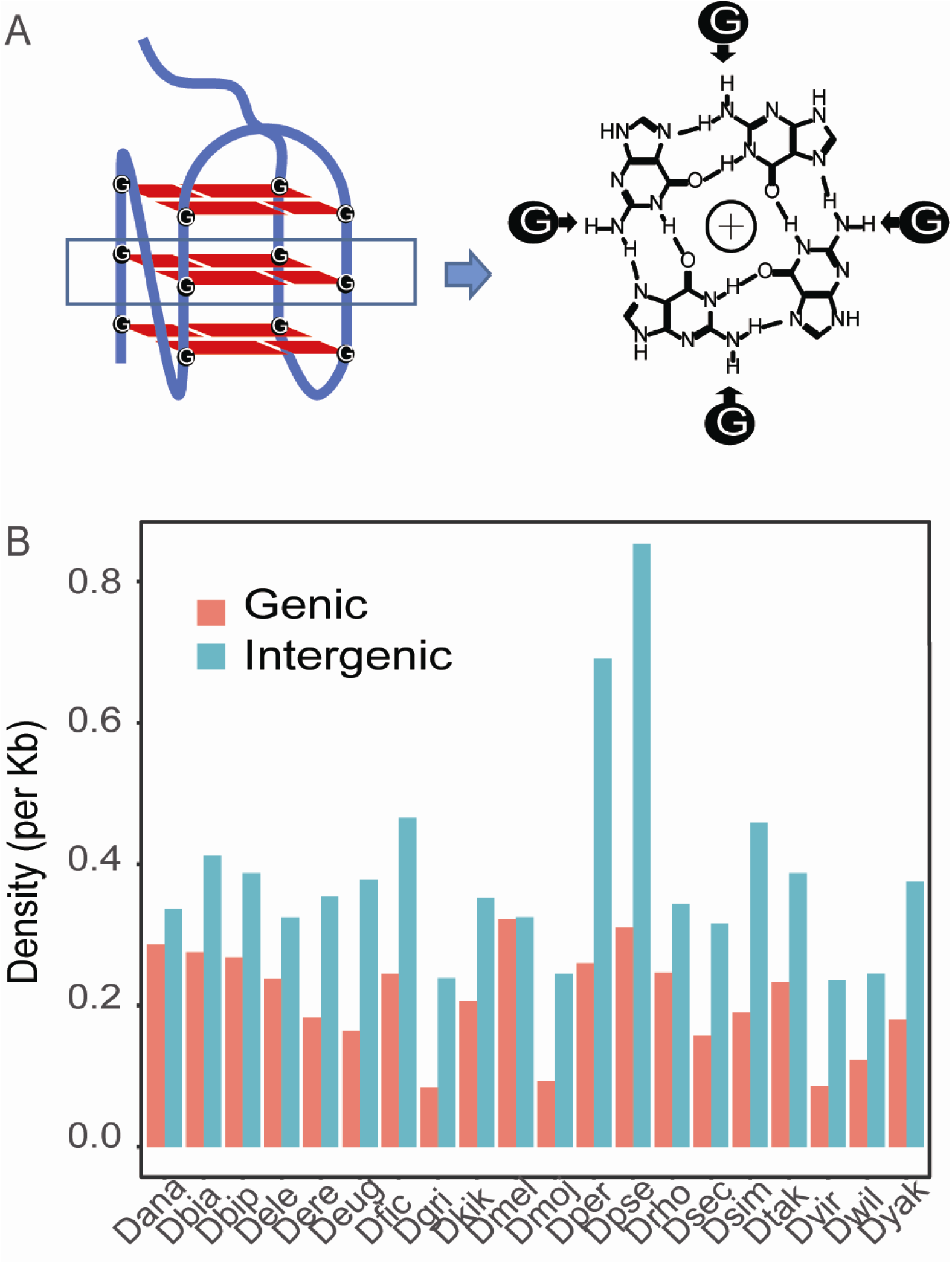
The enrichment of pGs in intergenic regions. (A) The structure of G-quadruplex. (B) The distribution of pG4s in the genome of 20 fly species. The G4s that have any overlap with gene region were referred as genic G4s, others were assigned as intergenic ones.

Considering repetitive regions encounter less selective constraints and nonfunctional elements were thus likely to accumulate there instead of non-repetitive regions, we classified intergenic pG4s into repetitive pG4s and non-repetitive pG4s based on whether or not they were overlapped with repetitive regions. We found that intergenic non-repetitive regions had the highest density with an average of 0.42/kb, followed by genic regions with an average of 0.32/kb and intergenic repetitive regions with an average of 0.23/kb (Table S1). This result suggested that intergenic pG4s, at least part of them, could be maintained as functional elements.

The hemizygosity of X chromosome in males led to different evolutionary forces acting on autosomes and X chromosome, followed by different distribution patterns of non-neutral elements in the genomes. We thus investigated the chromosomal distribution of pG4s to better understand why the genome, especially for intergenic regions, encodes so many of them. We found that pG4s were enriched on the X chromosome (2,552, 21.4 %) (Fig. 2A; P < 2.2 × 10-16, Fisher’s exact test). Such enrichment still existed even after the normalization for GC contents (Fig. S5). Interestingly, the pG4s on the X chromosome were longer (Fig. 2B; P < 2.2 × 10-16, Kruskal-Wallis test) and had higher quadruplex propensity than autosomal pG4s (Fig. 2C; P < 3.965 × 10^-13^, Kruskal-Wallis test), suggesting that pG4s, especially X-linked pG4s, were not neutral.

**Fig. 2.**
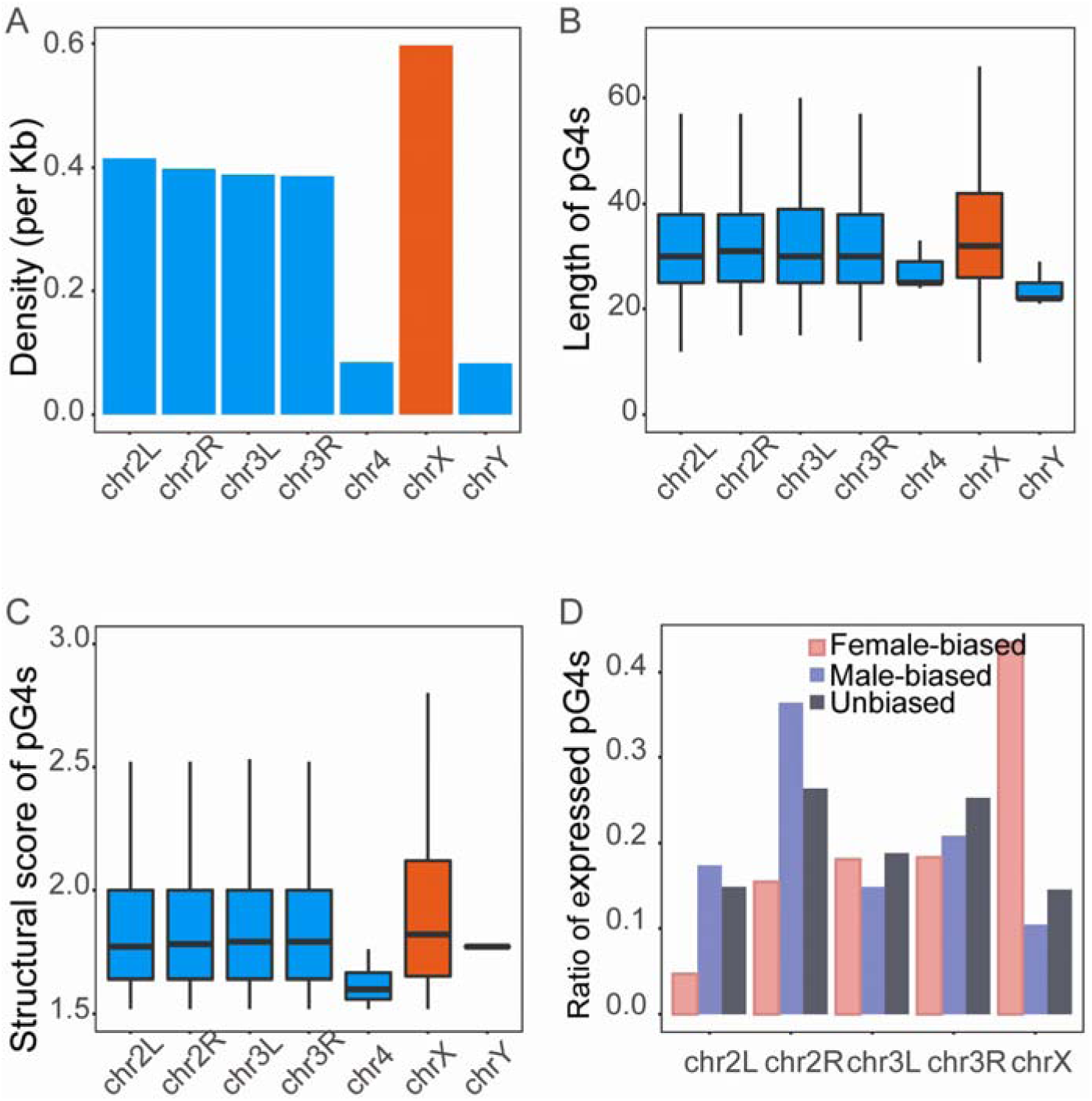
The best performance of X-linked pG4s in chromosomal distribution, length and structural stability relative to others. (A) The X chromosome has the highest density of pG4s than other chromosomes. (B) X-linked pG4s are longest than others. (C) The pG4s on the X chromosome are the most structurally stable than others. (D) The pG4s with male-biased expression were enriched on the X chromosome, while those with female-biased expression showed a deficiency on the X chromosome.

The distribution of sex-biased elements was shaped by various contrasting forces they were subjected to (*11, 12*). To find out some clues on the evolutionary mechanisms of pG4s based on the distribution of sex-biased pG4s, we classified all expressed pG4s into three groups, including male-biased, female-biased and unbiased pG4s. Among the expressed pG4s, 474 (7.56%) were male-biased expressed, 727 (11.60%) were female-biased expressed and 5,065 (80.83%) were unbiased. Compared with male-biased pG4s, female-biased pG4s had greater expression level (Fig. S6; P = 3.269 × 10^-5^, Wilcoxon test), quadruplex structural scores (Fig. S7; P = 1.823 × 10^-5^, Wilcoxon test) and length (Fig. S8; P = 1.675 × 10^-5^, Wilcoxon test), suggesting that they were maintained under selection.

We then investigated the distribution of these expressed pG4s across the chromosomes, and found that male-biased pG4s with expression were underrepresented on the X chromosome (P < 0.05, Wilcoxon), while female-biased pG4s were overrepresented on the X chromosome (Fig. 2D; P < 2.2 × 10^-16^, Wilcoxon test), consistent with the pattern of protein-coding genes (*13, 14*) and non-coding RNAs (*15*). The concurrent underrepresentation of X-linked male-biased pG4s and overrepresentation of female-biased pG4s could be explained by sexual antagonism for dominant genes (*16*) or the repression of dosage compensation (*17*) by G4s.

G4s were reported to suppress gene expression (*18, 19*). To test whether the enrichment of intergenic pG4s was responsible for the male-specific fine-tuning of X chromosome in dosage compensation, we then took advantage of RNA-seq data from fly tissues to measure the expression of pG4s. The pG4s with higher expression than 30% genic pG4s in any tissue were taken as expressed pG4s, while others were taken as unexpressed pG4s. In total, 21,009 of pG4s were expressed, including 2,739 (13.04%) intergenic pG4s. Expressed pG4s were underrepresented in intergenic regions. Moreover, we found lower expression level of intergenic pG4s than genic pG4s (Fig. S9; P < 2 × 10^-16^, Wilcoxon test), supporting that pG4s may suppress gene expression. We further tested whether the density of pG4s in a region was associated with its expression level with a sliding window strategy, and found windows with higher pG4 density had lower expression than those with lower pG4 density (Fig. 3A; P < 6.14 × 10^-14^, Wilcoxon test). The negative correlation between pG4 density and expression levels suggested the role of pG4s as suppressors. Next, we performed RNA-seq of *Drosophila* S2 cells treated with TMPyP4, a type of G-quadruplex stabilizer (Fig. 3B), and found the expression level of pG4s decreased with the increase of TMPyP4 concentration, supporting again the role of pG4s as suppressors.

**Fig. 3.**
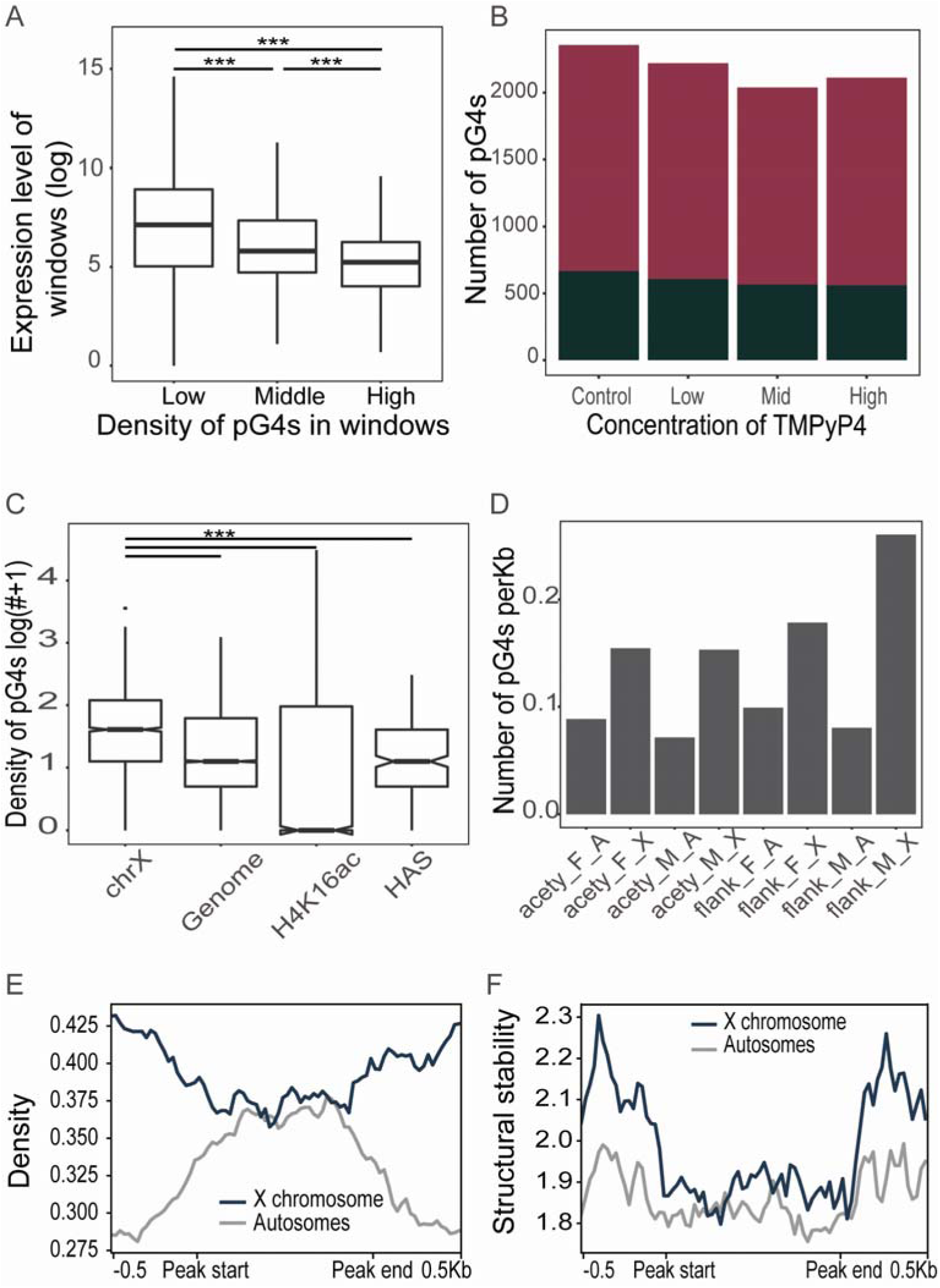
The low density and structural stability of G4s on the H4K16ac and HAS regions. (A) The correlation of a window between pG4 density and expression. A sliding window strategy was used to generate all genomic windows (1Kb) and divided into three groups based on thier pG4 density. We found the more pG4s density in a window, the less expression level the window showed. (B) The expression level of pG4s under different concentration of TMPyP4, DMSO was used as control. The pG4s show less expression level when were treated with TMPyP4 with higher concentration. (C) The pG4 density on the X chromosome, the whole genome, the H4K16ac regions and HAS regions. (D) X-linked specific flanking regions of H4K16ac in male showed the highest density of pG4s than other regions, acety means H4K16ac region, flank means flanking region of H4K16ac. (E and F) Density and structural stability profile of G4s in H4K16ac (scale to 1 kb) and flanking regions (upstream and downstream 500bp) on the male X chromosome and autosomes, respectively. Region from peak start to end indicated H4K16ac region.

The pG4s analyzed above were computationally predicted. To validate our observations, we thus reanalyzed the patterns of G4s identified with G4-seq in *Drosophila melanogaster (20)*. In total, 14,946 (32.5%, with K+ as stabilizing agent of G4-seq) and 17,495 pG4s (38.1%, with K+ and PDS as stabilizing agent of G4-seq) were supported by G4-seq (Table S2). The G4s identified by G4-seq on the X chromosome had higher density (∼1.9 fold), more stable structures, and greater length than those on the autosomes, consistent with observations from pG4s predicted in silico (Fig. S10-S12).

We further explored whether the enrichment of X-linked pG4s with decreased expression was associated with X-specific upregulation in males. We found the pG4 density at HAS (Genomic coordinates from (*3*)) on the X chromosome (0.263/kb) was not only lower than the average pG4 density on the X chromosome (0.488/kb), but also than that across the whole genome (0.335/kb). The pG4 density in H4K16ac regions on the X chromosome (0.357/kb) was lower than the average pG4 density on the X chromosome, although higher than that in H4K16ac regions of the whole genome (0.357/kb) (Fig. 3C). We further calculated the pG4 density in H4K16ac regions and their flanking regions on autosomes and the X chromosome in both sexes, and found highest pG4 density in the H4K16ac flanking regions on the male X chromosome (Fig. 3D). Peaks of both the density (Fig. 3E and Fig. S13) and the quadruplex propensity of pG4s (Fig. 3F and Fig. S14) were found at the boundaries of H4K16ac regions, suggesting that pG4s confined the acetylated regions mediated by MSL complex in dosage compensation.

To explore the evolutionary patterns of pG4, we further dated the pG4s based on their presence or absence in the phylogenetic tree of 20 *Drosophila* species (*21, 22*) (Fig. 4A). We found 3,328 (26.4%) pG4s were specific to *D.melanogaster*, and the correlation between age and pG4 number was −0.57 (Spearman correlation test). Moreover, the younger pG4s showed an excess on the X chromosome, while the older pG4s showed a paucity (Fig. S15), suggesting faster birth and death of X-linked pG4s than autosomal pG4s. We then separated pG4s into three groups (young, middle and old) based on their age, and observed that younger pG4s on the X chromosome were more likely to be female-biased (Fig 4B; P < 0.05, Fisher), suggesting that newly formed pG4s were more likely to repress dosage compensation. As a validation, we calculated the density of pG4s in *Drosophila miranda* and *Drosophila pseudoobscura* with a series of X chromosomes of diverse ages. The XL chromosomes of ∼60 million years old in their genomes were homologous to *D. melanogaster* X, and the XR chromosomes originated 15 million years ago, while the neo-X chromosome becoming a sex chromosome only 1 million years ago was specific to *D. miranda* (*23–25*). As expected, our result showed that the pG4 density on these sex chromosomes was higher than autosomes, and the pG4 density on the younger XR and neo-X were higher than that on the older XL (Fig. 4CD).

**Fig. 4.**
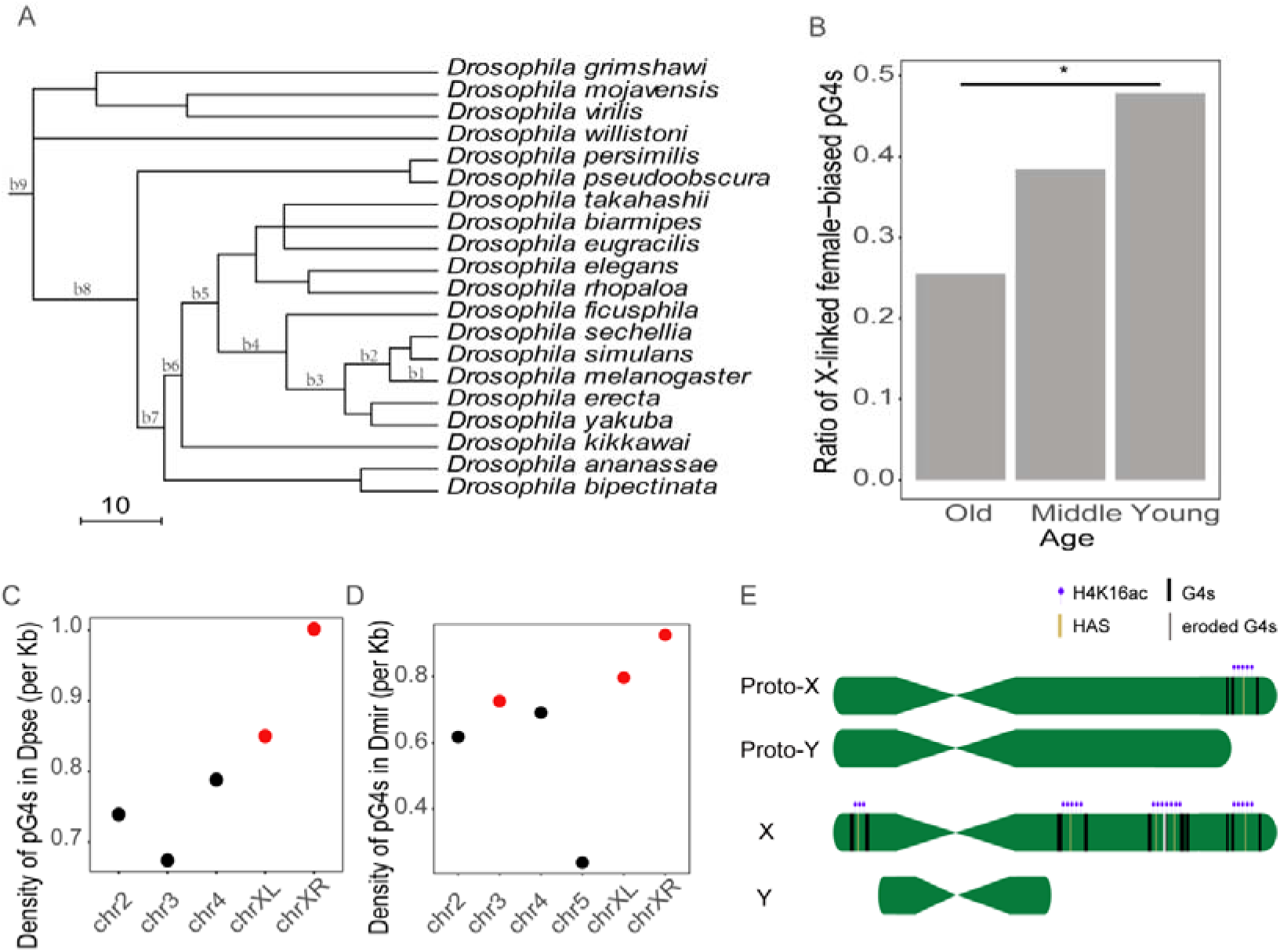
Gradually accumulated G4s on the X chromosome functions as insulators in dosage compensation. (A) Phylogenetic tree of 20 drosophila species. (B) X-linked G4s with female-biased expression were more likely to be younger. (C and D) Chromosomal density of pG4s in Dpse and Dmir. (E) Model of G4 evolution in dosage compensation. In the early stage of sex chromosome evolution, the Y chromosome still maintained abundant gene homologous to the X chromosome. Only the genes with single dose in male should be compensated, while other two-dose genes were not to be. The gradually formed G-quadruplexes enriched on the flanking regions around MSL binding sites and H4K16 acetylated regions, acting as insulators to precisely separate from the regions that should not be up-regulated. With the degradation of the Y chromosome, the X chromosome acquired more and more high-affinity sites (HAS) for large-scale dosage compensation, and the G4s located between two or many HAS may be eroded extend compensation regions.

A model was thus proposed for the evolution of G4s on the X chromosome (Fig. 4E). In the gradual degradation of Y chromosome, dosage compensation was required for one-dose proto-X-linked genes. To fine-tune the expression of these one-dose genes locally instead of chromosome-wide, G4s formed flanking a newly evolved HAS and acted as insulators to protect two-dose proto-X-linked regions from acetylation and upregulation. With the extension of one-dose proto-X regions, G4s were selected against to allow their upregulation.

## Acknowledgments

We thank Huijuan You, Dengguo Wei, Wenqiang Wu for providing TMPyP4, Zhihui Zhu for the gift of S2 cell. We also thank Manyuan Long, Brian Oliver, Jenny Graves, Charles Vinson, Xiao-Qin Xia, Yikang S. Rong, Qi Zhou, Jian Lu, Xionglei He, Jian-Quan Ni, Yu-Jie Fan and Wensheng Wei for helpful comments. Finally, I want to show my highest blessing to my father and mother, without them, there would be no me.

## Author contributions

S. Q. and Z.C. designed the project. S. Q. performed the analysis. L. C. carried out the experiment. S. Q. and Z. C. wrote the manuscript with all authors contributing to writing.

## Data and materials availability

Data for analyzing G-qudruplex expression alteration under different TMPyP4 concentration from this study have been submitted to the NCBI Gene Expression Omnibus (GEO; http://www.ncbi.nlm.nih.gov/geo/) under accession number GSE131691.

Supplementary Materials

Materials and Methods

Figures S1-S15

## Materials and Methods

### Genomes

Genome sequences of *Drosophila menalogaster* (dmel_r6.19_FB2017_06), *Drosophila pseudoobscura/* (Dpse_r3.04) were downloaded from flybase (http://flybase.org/). 18 genome sequences of D. *ana* (dana_caf1), D. *bip* (Dbip_2.0), D. *ere* (dere_caf1), D. *fic* (Dfic_2.0), D. *kik* (Dkik_2.0), D. *moj* (dmoj_caf1), D. *sec* (dsec_caf1), D. *tak* (Dtak_2.0), D. *wil* (dwil_caf1), D. *bia* (Dbia_2.0), D. *ele* (Dele_2.0), D. *eug* (Deug_2.0), D. *gri* (dgri_caf1), D. *per* (dper_caf1), D. *rho* (Drho_2.0), D. *sim* (ASM75419v2), D. *vir* (dvir_caf1), D. *yak* (dyak_caf1) were downloaded from National Center for Biotechnology Information (NCBI, https://www.ncbi.nlm.nih.gov/).

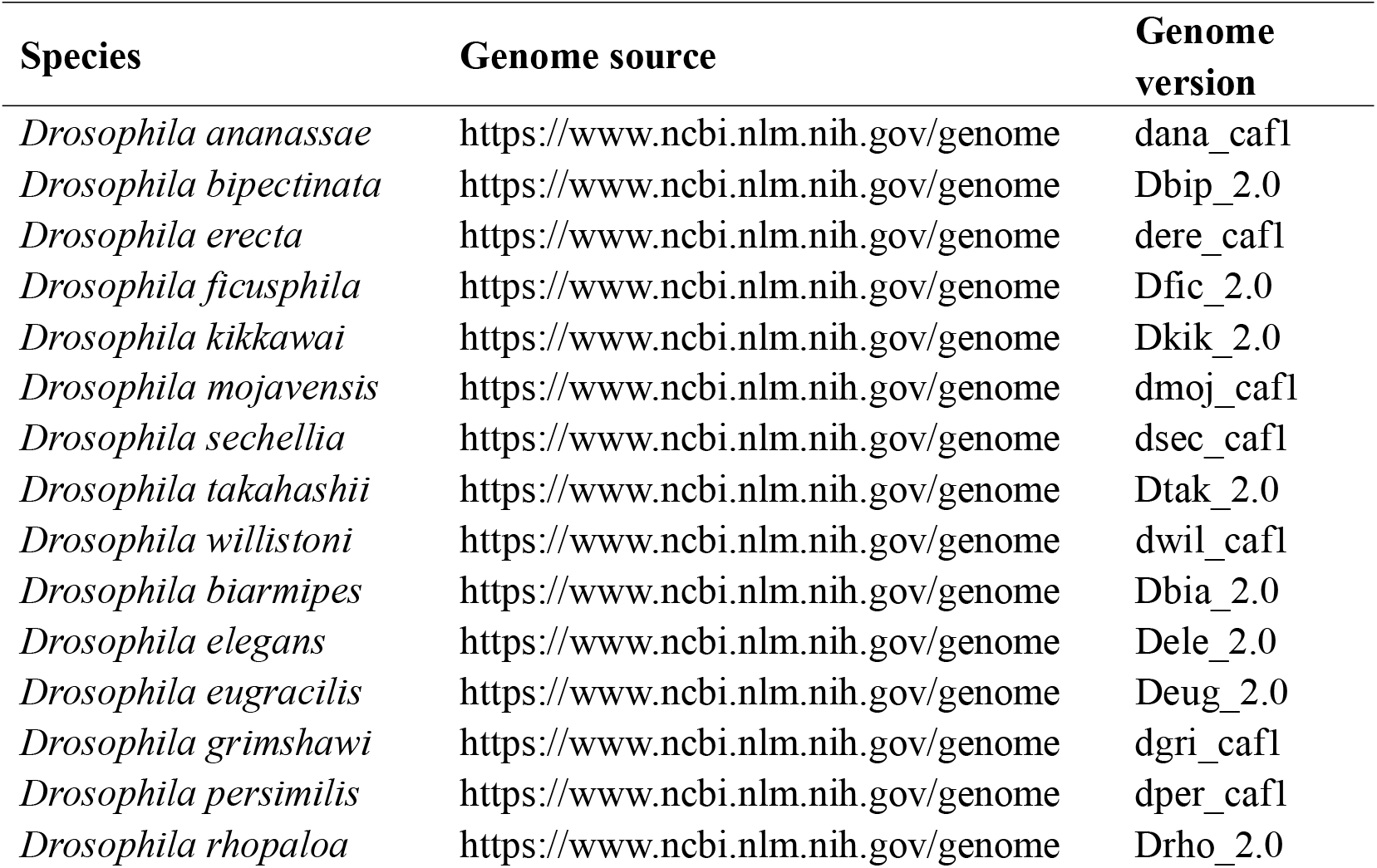
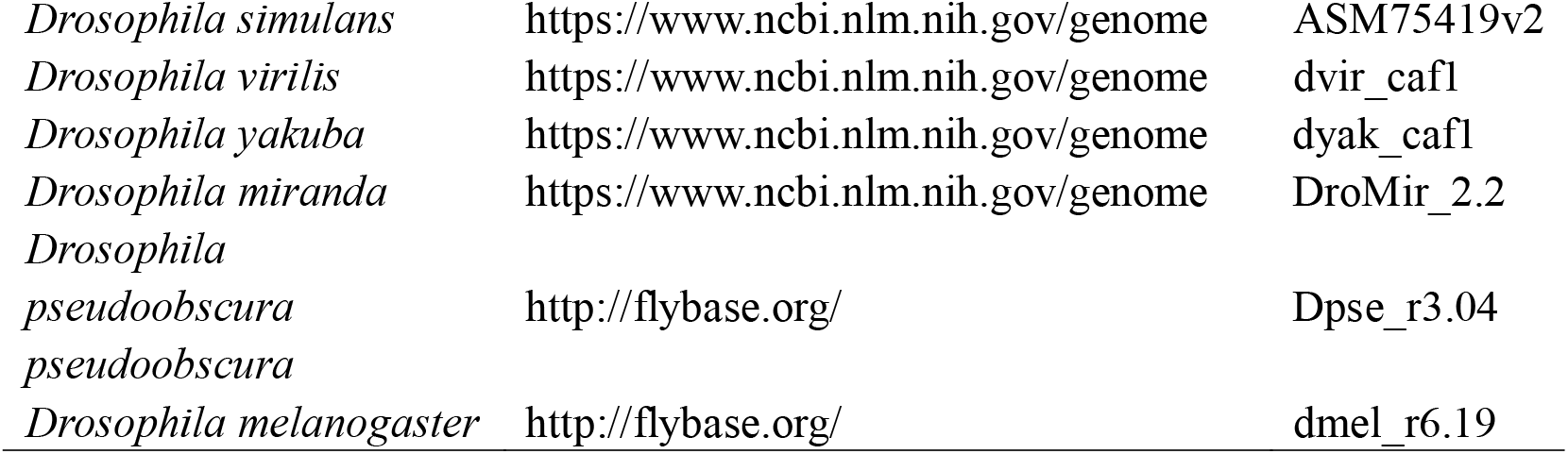

### Detecting pG4s in the fly genomes and simulated genome

We used R package G4Hunter to identify pG4s in Drosophila melanogaster genome (26). G4Hunter predicted pG4s using a score function in which positive scores were given for Guanine while negative scores were given for Cytosine in each strand of DNA sequences. To avoid overestimation of G4 number, overlapping pG4 sequences were fused. We specified cutoff as 1.50 to find pG4s with scores no less than 1.50. Meanwhile, we applied a sliding window strategy (window size: 5kb, step size: 2.5kb) to assess the correlation between number of pG4s and GC content. We searched pG4s on chromosome 2L, 2R, 3L, 3R, 4, X, and Y, and identified 45,943 pG4s in total (referred to as whole-genome pG4s).

Then we generated 1,000 artificial randomized fly genomes to assess statistical significance of our data. We randomly shuffled the genome while maintained their original DNA composition and length by shuffleseq, implemented in EMBOSS (v6.5.7) (32). Therefore, we compared the number of pG4s in simulated genome with that of in original genome but the identical nucleotide compositions and lengths.

### The classification of pG4s

We classified pG4s into repetitive and nonrepetitive PG4s based on repeats annotations produced by RepeatMasker (http://www.repeatmasker.org; last accessed Feb. 2, 2018) from UCSC Genome Browser. RepeatMasker classified low complexity regions and repeat regions into LTR, simple repeats, long Interspersed Elements (LINE), short Interspersed Elements (SINE) and other types of repeates. We referred the G4s having at least 1 nt overlap with any repeat as repetitive G4s and others as nonrepetitive G4s.

We also classified pG4s into genic and intergenic PG4s based on General Transfer Format (GTF) format gene annotation of D. melanogaster (Flybase r6.19) downloaded from Flybase (http://flybase.org/; last access Jan 27, 2018). We used the software bedtools command intersect (27) (v2.25.0) to search pG4s which had at least 1 nt overlap with any protein-coding genes, non-coding genes or psuedogenes and classified them into genic pG4s. Other pG4s were classified into intergenic pG4s. Furthermore, we separated pG4s into five groups based on their different length. The 20, 40, 60 and 80 quantiles of their length were 25, 28, 34 and 41. Thus pG4s were classified into five groups: SS, S, M, L and LL. GC contents in every region were calculated by bedtools nuc.

### Cell culture and treatment

Drosophila line 2 cells (S2) were maintained at 28 □ in Hyclone TNM-FH insect medium containing 10% Gibico serum and antibiotics (0.5 U/ml penicillin and 0.5 μg/ml streptomycin). Cells were treated with different concentration of TMPyP4 (25umol/L, 50umol/L, 100umol/L, respecticely), and DMSO (100μg/L) was severed as control. After 48 hours, the S2 cells were sampled for extraction of RNA.

### RNA extraction and sequencing

RNA samples from each individual were exacted using Trizol (from Invitrogen) according to the manufacturer’s protocol. Stranded PolyA+ RNA libraries were prepared with in-house kits (Majorbio). The library quality was assessed by checking the distribution of the fragment size using Agilent 2100 bioanalyzer (Agilent) and the quantity of the libraries were measured using qRT-PCR (TaqMan Probe). In total, 12 qualified strand-specific cDNA libraries were constructed and were sequenced on the Illumina HiSeq System.

### Expression profiling

RNA-seq data from fly somatic and gonadal tissues were downloaded from Gene Expression Omnibus (GEO) in NCBI (http://www.ncbi.nlm.nih.gov/geo/; accession numbers: GSE44612) (28). We then aligned sequence data, as well as raw reads in FASTQ format, by HISAT2 (v2.0.4) (29) to the reference genome (-x) and annotation (--known-splicesite-infile). To estimate the expression level of pG4s, we quantified counts of reads overlapping flanking 1 kb of each pG4 by slop and coverage implemented in bedtools, and normalized the read counts by R package DESeq2 (v1.18.1) (30). For each pG4, maximum normalized read counts among different tissues were regarded as expression level of pG4.

To define expression status of intergenic pG4s with high confidence, we used a strict cutoff derived from the expression level of genic pG4s. The median expression level of all genic pG4s was taken as the expression cutoff for intergenic pG4s. Then intergenic pG4s with normalized read count higher than the cutoff were defined as expressed pG4s, and others were unexpressed.

### Dating G-quadruplexes on Phylogenetic Tree of Drosophila

To date all pG4s on Drosophila phylogenetic tree, we applied a modified pipeline based on previous method (21, 22). We performed whole-genome pairwise alignments between D. melanogaster and other 19 Drosophila species (D.ananassae, D.biarmipes, D.bipectinata, D.elegans, D.eugracilis, D.ficusphila, D.kikkawai, D.mojavensis, D.pseudoobscura, D.rhopaloa, D.simulans, D.takahashii, D.virilis, D.yakuba, D.erecta, D.sechellia, D.willistoni, D.grimshawi and D.persimilis). Based on UCSC genome browser mannual, we generated reciprocal best LiftOver chain files from the alignments and aligned D. melanogaster to other Drosophila species. We defined a pG4 as present in a species if it could be projected to the species no matter whether it was still a pG4s motif in the species. Finally, we dated all pG4s on the phylogenetic tree and assigned pG4s into different evolutionary branches. The pG4s assigned to branch1-3 were classified as young, those in branch4-6 were classified as middle and the rest were regarded as old.

### MSL ChIP-seq and HAS analysis

ChIP-seq data for H4K16ac analysis were downloaded from NCBI under accession number GSE109901. We aligned the sequence data by bowtie2 (31) to the reference genome with default parameter. BAM files from ChIP-seq were used for peak calling with MACS2 (32). A bed file containing genomic coordinates of HAS was downloaded from (3) and converted to corresponding version by UCSC liftover with default parameter.

Profile of G4 in H4K16ac (scale to 1 kb) and flanking regions (500 bp) was analyzed by deeptools.

