## Supplemental Figures for "Enriched G-quadruplexes on the *Drosophila* Male X Chromosome Function as Insulators of Dosage Compensation Complex"

Supplementary Figures

Fig. S1.

**The number of pG4s in 20 fly species.** Three cutoffs of G4hunter (1.0, 1.2, 1.5) were used. The red, green and blue column indicates the loose, middle and strict cutoff, respectively.

Fig. S2.

**The number of pG4s in simulated fly genomes.** We randomly shuffled the D. melanogaster genome 1000 times to generated simulated genomes while maintained their original DNA composition and length, and then identified pG4s with the same criteria.

Fig. S3.

**The relationship between the number of pG4s and GC content in a window.** A sliding window strategy (size 5kb, step size 2.5kb) was used to evaluate the relationship between them. Each point represents a window. There was a positive correlation between number of pG4s and GC content. A pG4 of which whole falls into a window was assigned to belong to this window. There was a positive correlation between number of pG4s and GC content.

Fig. S4.

**The GC content of genic and intergenic regions across 20 *Drosophila* species.** The red column indicates GC contents of genic regions, and the blue column indicates GC contents of intergenic regions. The GC contents of genic regions are higher than that of intergenic regions in 20 drosophila species.

Fig. S5.

**The density of pG4s on each chromosome after correlation of GC content.** The density of pG4s on all chromosomes was calculated after correlation of GC content as the density of pG4s is related to the GC content. The X chromosome has the highest density of pG4s than others (Fig. 2A), such enrichment still exists after correlation of GC content.

Fig. S6.

**The expression level of pG4s with different sex biased expression.** The pG4s with female-biased expression have the higher expression level than both male-biased and unbiased ones.

Fig. S7.

**The quadruplex propensity of pG4s with different sex biased expression.** The pG4s with female-biased expression have the higher quadruplex propensity than both male-biased and unbiased ones.

Fig. S8.

**The Length of pG4s with different sex biased expression.** The pG4s with female-biased expression are longer than both male-biased and unbiased ones.

Fig. S9.

**The expression levels of pG4s in genic regions and intergenic regions.** The expression level of pG4s in genic regions was higher than that in intergenic regions.

Fig. S10.

**Normalized density of G4s on chromosomes under different treatments.** Density of G4s on chromosomes normalized for GC content under K (A) and K+PDS (B), respectively.

Fig. S11.

**Structural stability of G4s on chromosomes under different treatments.** Structural stability of G4s on chromosomes under K (A) and K+PDS (B), respectively.

Fig. S12.

**Length of G4s on chromosomes under different treatments.** Length of G4s on chromosomes under K (A) and K+PDS (B), respectively.

Fig. S13.

**Density profile of G4s in the H4K16ac and flanking regions on the male X chromosome and autosomes, respectively, under K (A) and K+PDS (B).** The black line indicates the density in the X chromosome and the grey line indicates the density on the autosomes. The density of G4s in the H4K16ac regions on the X chromosome is lower than the average density of G4s on the X chromosome under both K treatment and K+PDS treatment, even lower than the average density of G4s on the autosomes under K treament, while the density of G4s on flanking regions on the X chromosome is higher than the average density on the X chromosome.

Fig. S14.

**Structural stability profile of G4s in the H4K16ac and flanking regions on the male X chromosome and autosomes, respectively, under K (A) and K+PDS (B).** The black line indicates the density in the X chromosome and the grey line indicates the density on the autosomes.

Fig. S15.

**The ratio of pG4s with different ages.** The columns with different color indicate the number of pG4s with different ages.

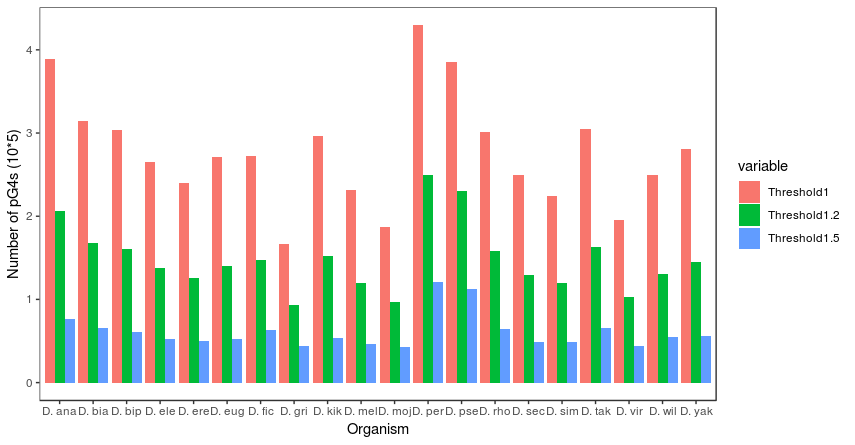

**Fig. S1.**

The number of pG4s in 20 fly species. Three cutoffs of G4hunter (1.0, 1.2, 1.5) were used. The red, green and blue column indicates the loose, middle and strict cutoff, respectively.

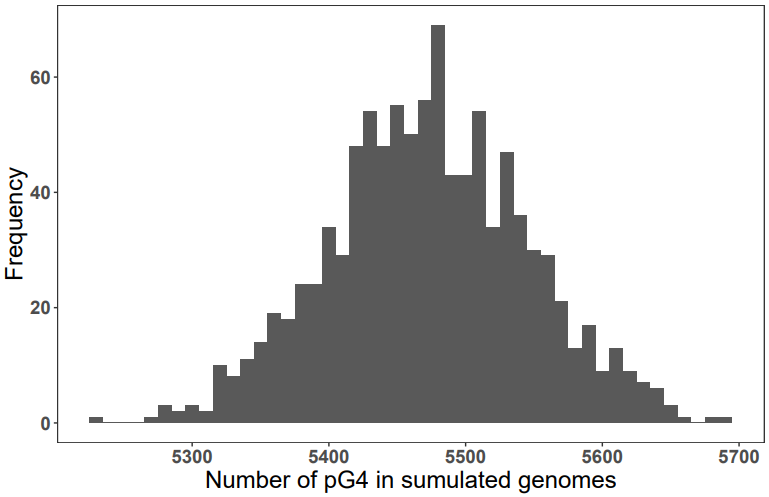

**Fig. S2.**

The number distribution of pG4s in simulated genomes. We randomly shuffled the *D. melanogaster* genome 1000 times to generated simulated genomes while maintained their original DNA composition and length, and then identified pG4s with the same criteria.

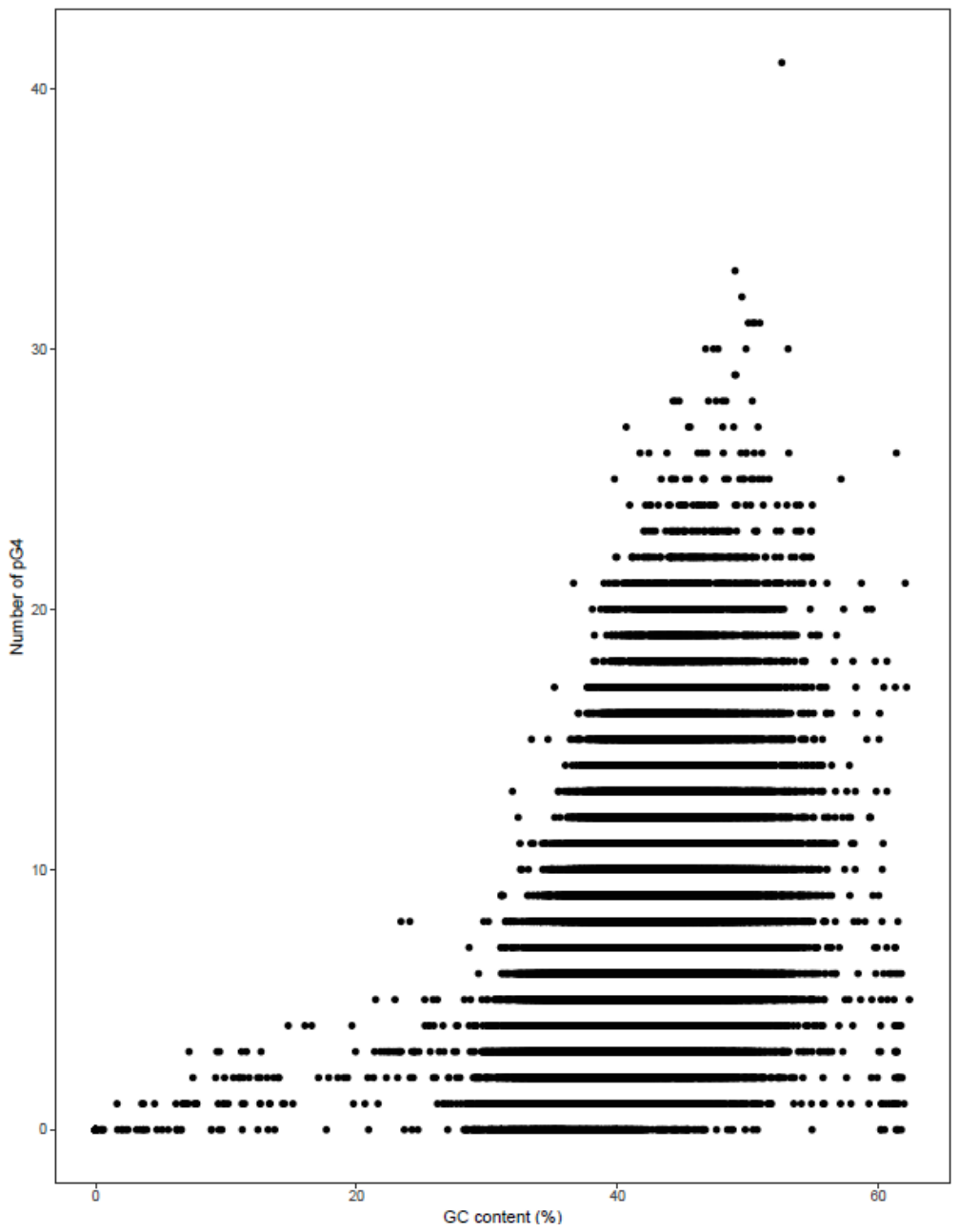

**Fig. S3.**

The relationship between the number of pG4s and GC content in a window. A sliding window strategy (size 5kb, step size 2.5kb) was used to evaluate the relationship between them. Each point represents a window. There was a positive correlation between number of pG4s and GC content. A pG4 of which whole falls into a window was assigned to belong to this window. There was a positive correlation between number of pG4s and GC content.

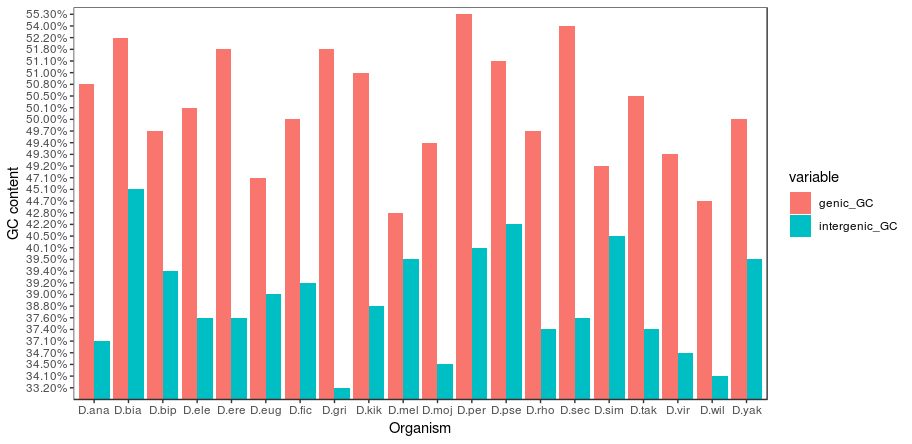

**Fig. S4.**

The GC contents of genic and intergenic regions across 20 *Drosophila* species. The red column indicates GC contents of genic regions, and the blue column indicates GC contents of intergenic regions. The GC contents of genic regions are higher than that of intergenic regions in 20 drosophila species.

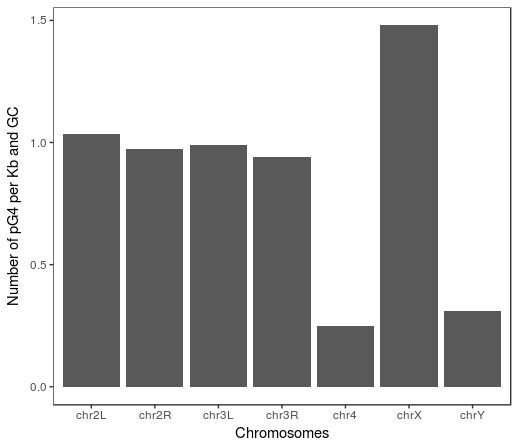

**Fig. S5.**

The density of pG4s on each chromosome after the normalization for GC content. The density of pG4s on all chromosomes was calculated after correlation of GC content as the density of pG4s is related to the GC content. The X chromosome has the highest density of pG4s than others (Fig. 2A), such enrichment still exists after correlation of GC content.

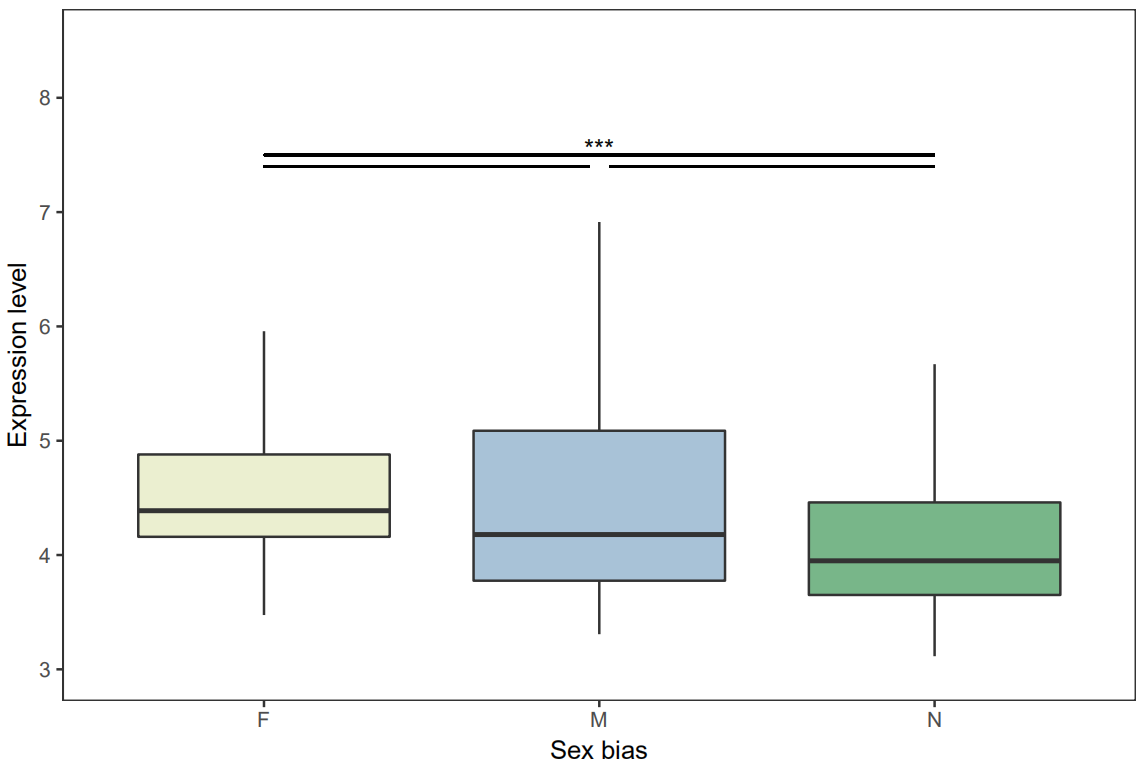

**Fig. S6.**

The expression level of pG4s with different sex biased expression. The pG4s with female-biased expression have the higher expression level than both male-biased and unbiased ones.

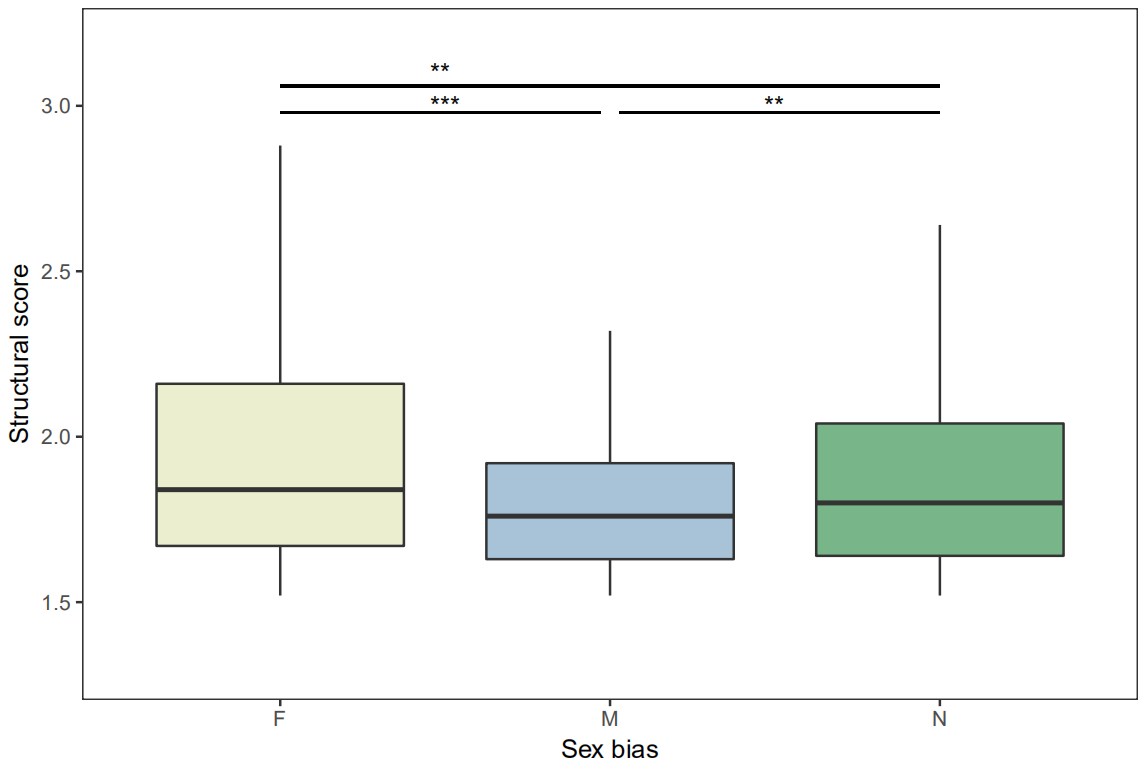

**Fig. S7.**

The quadruplex propensity of pG4s with different sex biased expression. The pG4s with female-biased expression have the higher quadruplex propensity than both male-biased and unbiased ones.

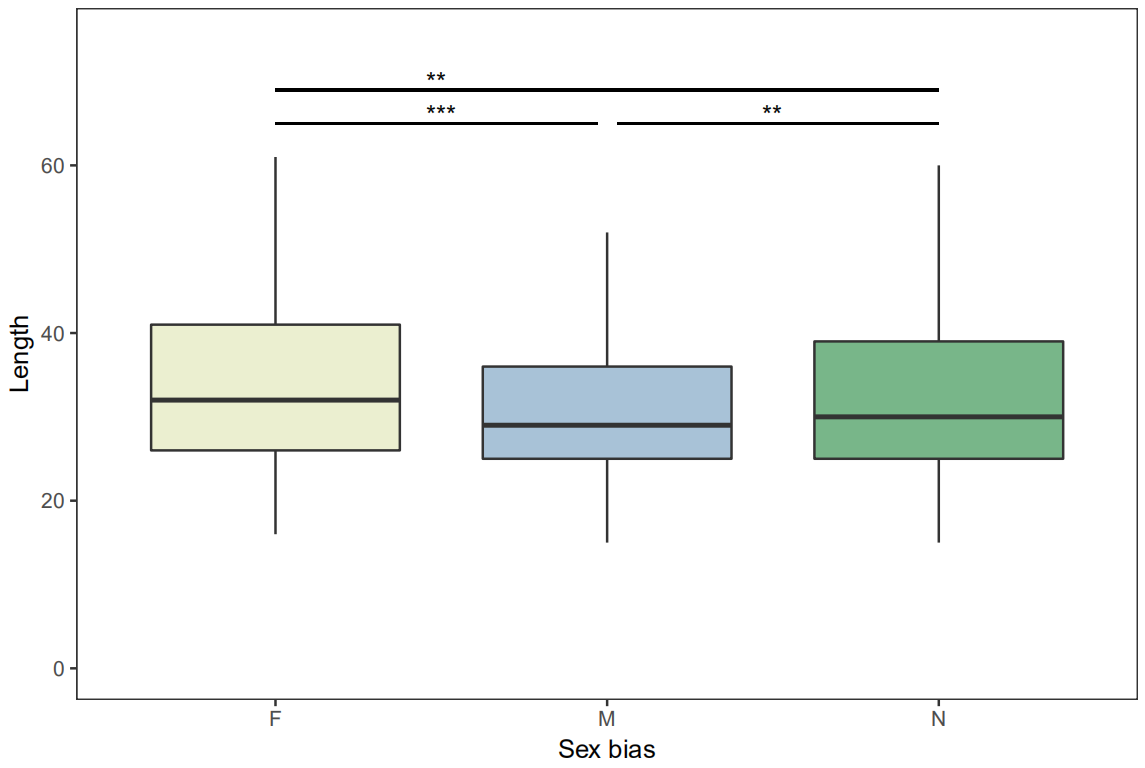

**Fig. S8.**

The length of pG4s with different sex biased expression. The pG4s with female-biased expression are longer than both male-biased and unbiased ones.

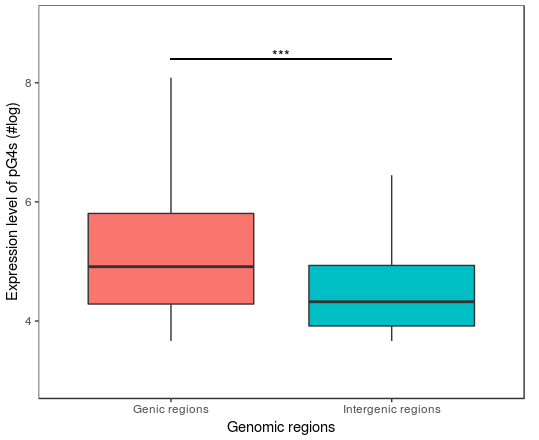

**Fig. S9.**

The expression levels of pG4s in genic regions and intergenic regions. The expression level of pG4s in genic regions was higher than that in intergenic regions.

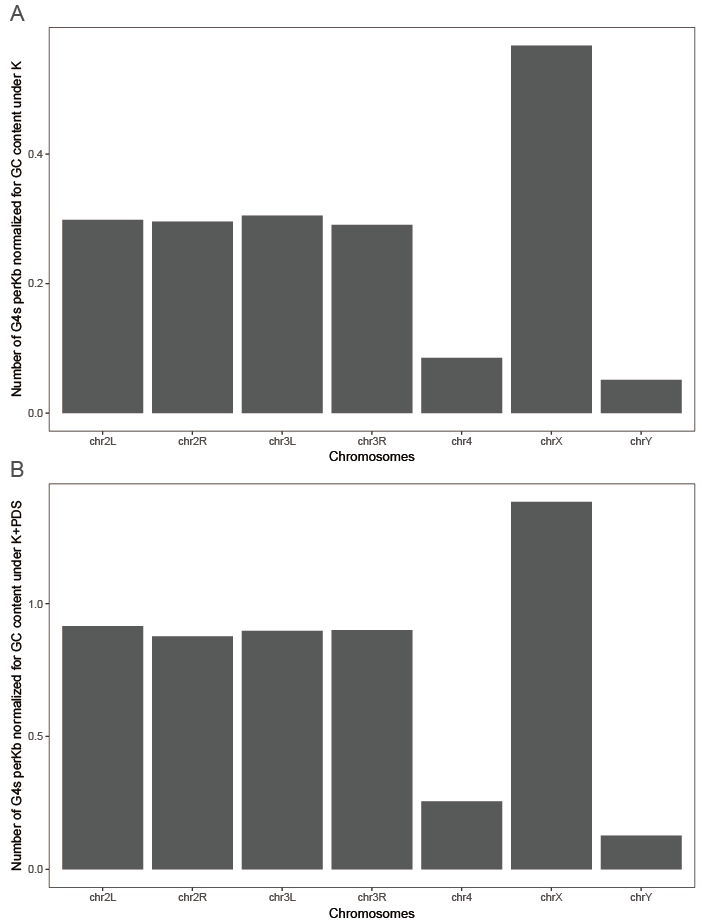

**Fig.S10**

Normalized density of G4s on chromosomes under different treatments. Density of G4s on chromosomes normalized for GC content under K (A) and K+PDS (B), respectively.

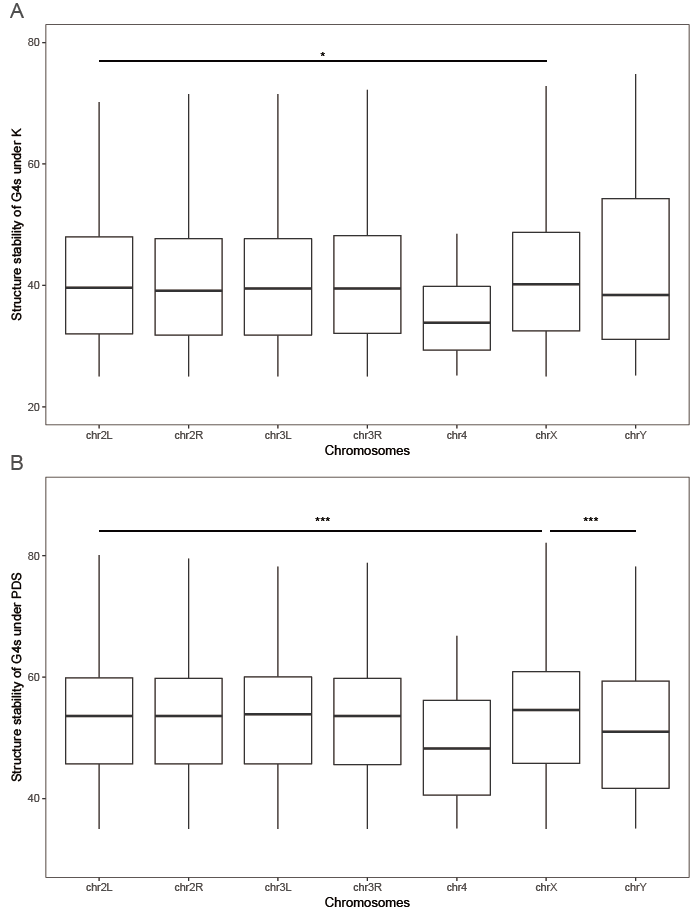

**Fig. S11**

Structural stability of G4s on chromosomes under different treatments. Structural stability of G4s on chromosomes under K (A) and K+PDS (B), respectively.

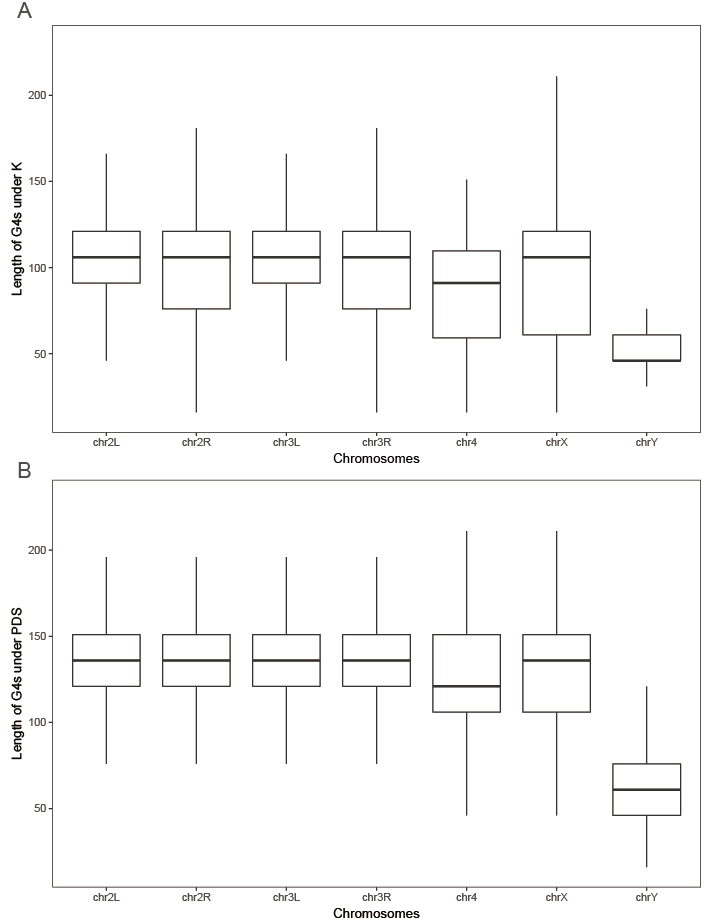

**Fig. S12**

Length of G4s on chromosomes under different treatments. Length of G4s on chromosomes under K (A) and K+PDS (B), respectively.

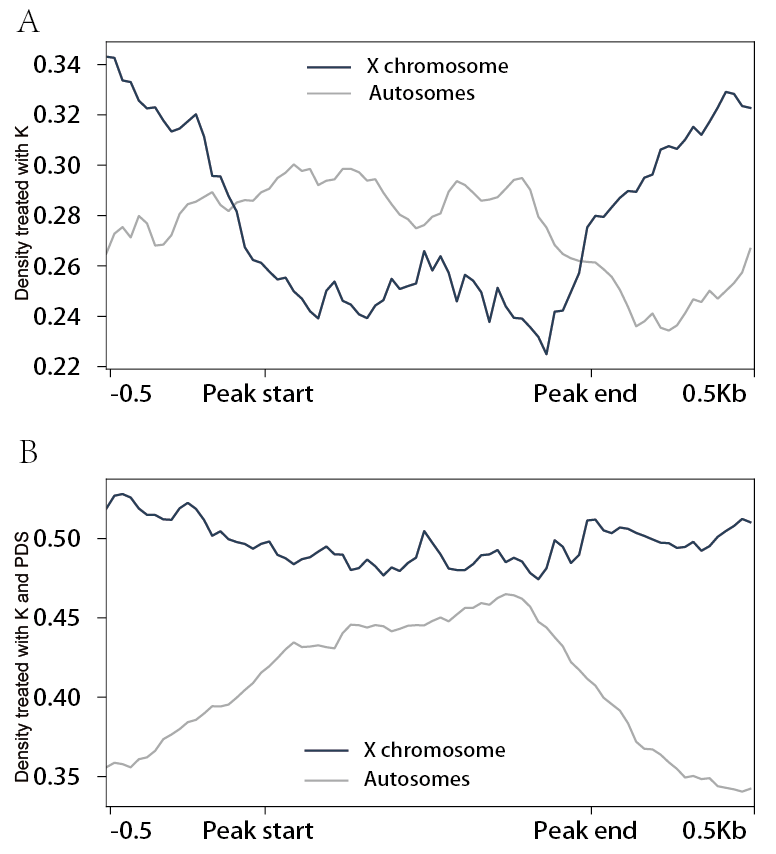

**Fig. S13.**

Density profile of G4s in the H4K16ac and flanking regions on the male X chromosome and autosomes, respectively, under K (A) and K+PDS (B). The black line indicates the density in the X chromosome and the grey line indicates the density on the autosomes. The density of G4s in the H4K16ac regions on the X chromosome is lower than the average density of G4s on the X chromosome under both K treatment and K+PDS treatment, even lower than the average density of G4s on the autosomes under K treament, while the density of G4s on flanking regions on the X chromosome is higher than the average density on the X chromosome.

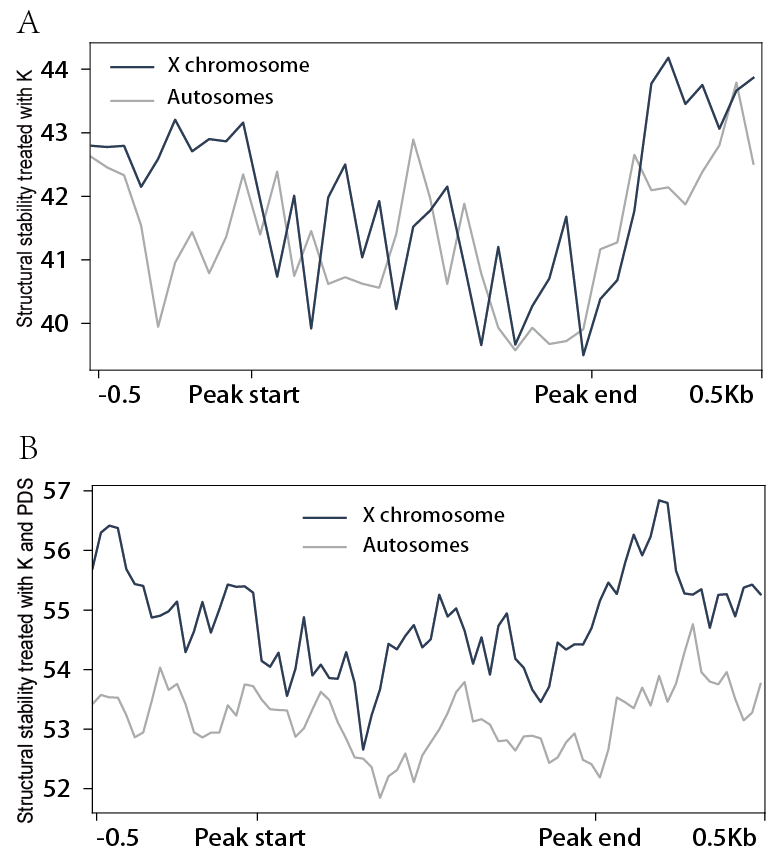

**Fig. S14.**

Structural stability profile of G4s in the H4K16ac and flanking regions on the male X chromosome and autosomes, respectively, under K (A) and K+PDS (B). The black line indicates the density in the X chromosome and the grey line indicates the density on the autosomes.

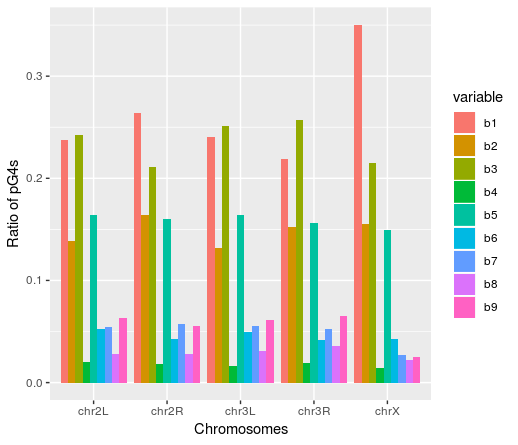

**Fig. S15.**

The ratio of pG4s with different ages. The columns with different color indicate the number of pG4s with different ages.
